# Incomplete penetrance for isolated congenital asplenia in humans with mutations in translated and untranslated *RPSA* exons

**DOI:** 10.1101/356832

**Authors:** Alexandre Bolze, Bertrand Boisson, Barbara Bosch, Alexander Antipenko, Matthieu Bouaziz, Paul Sackstein, Malik Chaker-Margot, Vincent Barlogis, Tracy Briggs, Elena Colino, Aurora C. Elmore, Alain Fischer, Ferah Genel, Angela Hewlett, Maher Jedidi, Jadranka Kelecic, Renate Krüger, Cheng-Lung Ku, Dinakantha Kumararatne, Sam Loughlin, Alain Lefevre-Utile, Nizar Mahlaoui, Susanne Markus, Juan-Miguel Garcia, Mathilde Nizon, Matias Oleastro, Malgorzata Pac, Capucine Picard, Andrew J. Pollard, Carlos Rodriguez-Gallego, Caroline Thomas, Horst Von Bernuth, Austen Worth, Isabelle Meyts, Maurizio Risolino, Licia Selleri, Anne Puel, Sebastian Klinge, Laurent Abel, Jean-Laurent Casanova

## Abstract

Isolated congenital asplenia (ICA) is the only known human developmental defect exclusively affecting a lymphoid organ. In 2013, we showed that private deleterious mutations in the protein-coding region of *RPSA*, encoding ribosomal protein SA, caused ICA by haploinsufficiency with complete penetrance. We reported seven heterozygous protein-coding mutations in 8 of the 23 kindreds studied, including 6 of the 8 multiplex kindreds. We have since enrolled 33 new kindreds, 5 of which are multiplex. We describe here eleven new heterozygous ICA-causing *RPSA* protein-coding mutations, and the first two mutations in the 5’-UTR of this gene, which disrupt mRNA splicing. Overall, 40 of the 73 ICA patients (55%) and 23 of the 56 kindreds (41%) carry mutations located in translated or untranslated exons of *RPSA*. Eleven of the 43 kindreds affected by sporadic disease (26%) carry *RPSA* mutations, whereas 12 of the 13 multiplex kindreds (92%) carry *RPSA* mutations. We also report that six of eighteen (33%) protein-coding mutations and the two (100%) 5’-UTR mutations display incomplete penetrance. Three mutations were identified in 2 independent kindreds, due to a hotspot or a founder effect. Lastly, RPSA ICA-causing mutations were demonstrated to be *de novo* in 7 of the 23 probands. Mutations in *RPSA* exons can affect the translated or untranslated regions and can underlie ICA with complete or incomplete penetrance.

## Introduction

Isolated congenital asplenia (ICA, MIM271400) is characterized by the absence of a spleen at birth in humans without other developmental defects. It renders otherwise healthy children susceptible to life-threatening invasive infections with encapsulated bacteria, typically *Streptococcus pneumoniae*, but occasionally *Neisseria meningitidis* and *Haemophilus influenzae* b (1, 2). Asplenia can be detected by ultrasound (US) or computed tomography (CT) scans of the abdomen. The associated defect of spleen phagocytic function is confirmed by the detection of Howell-Jolly bodies on a blood smear. ICA is the only known developmental defect of humans restricted exclusively to a lymphoid organ, as the DiGeorge (3) and Nude (4) syndromes, for example, involve both the thymus and other tissues. A retrospective study in France showed that this condition affects at least 0.51 per 1 million newborns per year (2). However, the incidence of ICA is probably higher, as individuals with ICA may not present their first severe infection until adulthood (5), and may be incidentally diagnosed with ICA in the absence of infection (6–9). Most cases of ICA are sporadic, but multiplex kindreds exist, and the main mode of inheritance of ICA seems to be autosomal dominant (AD).

In 2013, we tested the hypothesis of genetic homogeneity underlying ICA in at least some unrelated patients, by looking for rare nonsynonymous variants of the same gene in several patients from different kindreds. Using whole-exome sequencing (WES), we identified seven heterozygous mutations of *RPSA* in 19 patients from 8 kindreds, among 36 patients from 23 kindreds studied in total (5). This includes individuals from these kindreds for whom we collected DNA after the publication of our original study. The mutations were located in protein-coding regions and included one frameshift duplication (p.P199Sfs*25), one nonsense (p.Q9*), and five missense (p.T54N, p.L58F, p.R180W, p.R180G, p.R186C) mutations. Mutations of *RPSA* were more frequent in familial than in sporadic cases, being detected in 6 of the 8 multiplex kindreds (75%) and 2 of the 15 kindreds with sporadic disease (13%). All mutations were private to the ICA cohort, three occurred *de novo* (p.T54N, p.L58F, and p.R180W), and one (p.R180G) was recurrent, due to a mutational hotspot rather than a founder effect. Complete penetrance was observed in all 8 kindreds, as all 19 individuals carrying a rare heterozygous nonsynonymous mutation of *RPSA* had ICA. It should be noted that we did not investigate whether synonymous or non-protein-coding mutations in *RPSA* exons could cause ICA in our previous paper. Moreover, exon 1 of *RPSA*, which encodes only the 5’-UTR of *RPSA*, and the part of exon 7 encoding the 3’-UTR were not covered by the exome capture kit used in our previous study.

Since our first description of *RPSA* mutations in 2013, we have enrolled 37 additional ICA patients from 33 new and independent kindreds. Nine of these 33 kindreds approached us spontaneously after reading about our research online. Our international ICA cohort now comprises 73 patients from 56 kindreds with diverse ancestries and living on four continents (**Figure S1**). Patients were included if they had asplenia or a severely hypoplastic spleen documented by US, CT scan, or autopsy (10) (**Table S1**, **Figure S2**), and excluded if they had other developmental defects, such as an associated congenital heart malformation (a type of heterotaxy known as Ivemark or asplenia syndrome) (OMIM # 208530). The congenital nature of asplenia is documented at birth in rare cases, occasionally inferred from family history in multiplex kindreds, but is typically suspected in index cases after an episode of invasive infection (**Table S1**). The protein-coding region of *RPSA* was analyzed by Sanger sequencing in the 37 newly recruited patients, and we searched for *RPSA* copy number variation by multiplex ligation-dependent probe amplification (MLPA) in kindreds with no mutations in the protein-coding regions of *RPSA*. We also tested the hypothesis that non-coding mutations of *RPSA* can underlie ICA. Sanger sequencing was, therefore, performed on exon 1 and the flanking intronic regions (the word ‘exon’ will hereafter refer to the exon as well as the intronic bases at the intron-exon or exon-intron junctions), and the non-protein-coding parts of exons 2 and 7, which encode the 5’-and 3’-UTR of *RPSA*.

## Results

### A model for the genetic architecture of ICA caused by *RPSA* mutations

The field of human genetics has benefited from the recent release of large databases reporting allele frequencies for variants observed in the exomes (123,000) and genomes (75,000) of about 200,000 individuals (11). These new tools can be used to rule variants out as the cause of a disease on the basis of their allele frequency in the general population. However, several parameters (prevalence, inheritance, penetrance, genetic and allelic heterogeneity) must be taken into account before defining the highest allele frequency in these databases considered plausible for an ICA-causing variant. Based on an estimate of about one in a million children being admitted to hospital for ICA (2), and the fact that about 40% of individuals with ICA were never admitted to hospital during childhood (10 of the 25 patients that were not on prophylactic antibiotics during childhood, in our cohort) (**Figure 1A**, **Table S1**) (5), we estimate the global prevalence of ICA to be about 1 in 600,000. We found mutations in the protein-coding sequence of *RPSA* to display complete penetrance for ICA in our previous study, but AD disorders by haploinsufficiency generally display incomplete penetrance (12, 13). We therefore chose to apply a model of high penetrance but not complete to be conservative and selected 75% penetrance. As variants of *RPSA* were shown to be the genetic cause of ICA in eight of 23 kindreds in our previous study, we estimated the *RPSA* gene to be responsible for about 30% of ICA cases (genetic heterogeneity of 0.3). We observed the same mutation twice in the eight ICA kindreds with *RPSA* mutations. We therefore hypothesized that a single mutation could explain up to 25% of the ICA kindreds (allelic heterogeneity of 0.25). Using these estimates, we were able to calculate a maximum plausible allele frequency for an ICA-causing variant in a monoallelic dominant model, as follows: 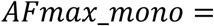 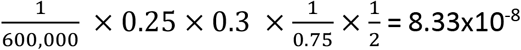 (14). Therefore, in our monoallelic model, the maximum tolerated allele count for an ICA-causing variant in the two combined and complete databases (~200,000 individuals) would be 0 (95% confidence interval). The maximum tolerated allele count for an ICA-causing variant in these databases restricted to WGS (75,000 individuals) would also be 0. The seven nonsynonymous mutations identified in our 2013 study (5) are absent from the gnomAD and Bravo databases (accessed in February 2018), and, therefore, meet these criteria.

**Figure 1:**
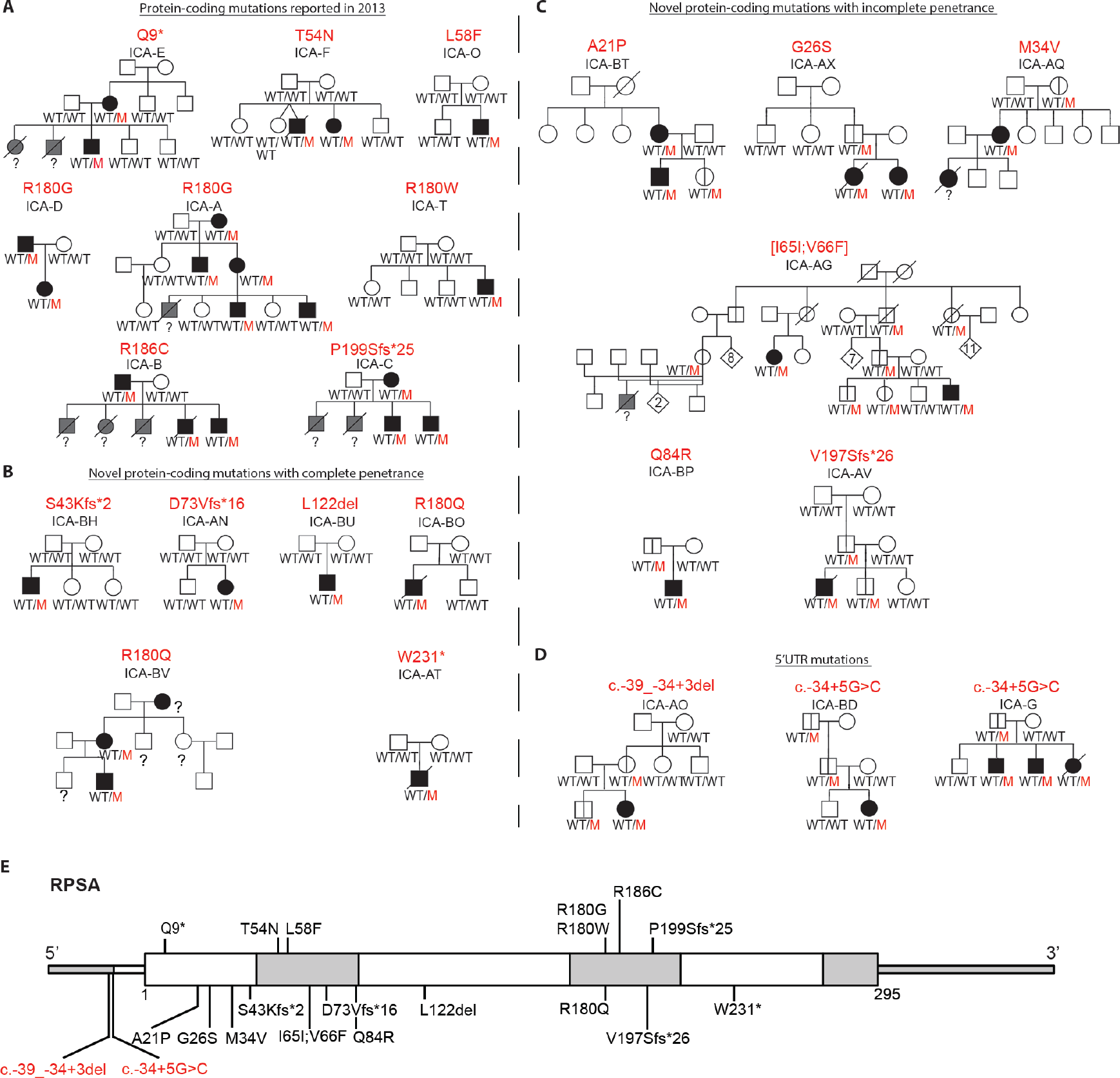
Overview of the 23 ICA kindreds in our cohort who carry mutations in *RPSA*. A. Eight kindreds were reported in our initial publication: all had non-synonymous and fully penetrant mutations for ICA. **B. and C.** Twelve newly recruited kindreds had novel non-synonymous mutations, displaying complete (B) or incomplete penetrance (C) for ICA. **D.** Three ICA kindreds with mutations in the 5’-UTR of *RPSA*. **E.** Schematic view of all of the ICA-causing mutations in *RPSA*. Mutations reported in our initial study are indicated above the gene schema. New mutations identified and reported in this study are indicated underneath the gene schema. Alternate white and grey color indicate the 7 different exons from the canonical transcript of the gene (ENST00000301821).

### Identification of previously unknown protein-coding mutations in *RPSA*

We therefore hypothesized that nonsynonymous variants of *RPSA* absent from gnomAD and Bravo would be present in about a third of the newly recruited kindreds (**Figure S1**). Sanger sequencing of the protein-coding region of *RPSA* was performed for the 33 newly recruited kindreds. We identified eleven heterozygous coding mutations, in 12 kindreds, none of which was previously reported in ICA patients. None of these mutations are present in any of the public databases, or in our in-house cohort of 4,500 exomes from patients with various infectious diseases (as of February 2018). The mutations can be grouped as (i) six missense mutations (p.A21P, p.G26S, p.M34V, p.[I65I;V66F], p.Q84R, p.R180Q), (ii) one in-frame deletion (p.L122del), (iii) three small frameshift insertions or deletions (p.S43Kfs*2, p.D73Vfs*16, and p.V197Sfs*26), and (iv) one nonsense mutation (p.W231*) (**Figure 1B, 1C, 1E**). All known ICA patients in the new 5 multiplex families for whom DNA was available carried a missense mutation of *RPSA* (**Figure 1B, 1C**). No gDNA was available for one of the deceased ICA patients from family ICA-AQ, for another deceased individual who probably had ICA from family ICA-AG, and for one ICA patient from family ICA-BV who was recently recruited to participate in the study. The mutations in kindreds ICA-AN, ICA-AT, ICA-BH and ICA-BU were found to have occurred *de novo*, after microsatellite analysis (MSA) to validate parent-child relationships of the DNA samples we sequenced. The mutation observed in ICA-AV was found to have occurred *de novo* in the father of the index case (first-generation relative) (**Figure S3**). Mutation p.R180Q is recurring in 2 families, but we were not able to analyze the full haplotype around the mutation in family ICA-BO. Finally, we performed MLPA in the remaining kindreds, to detect copy number variations overlapping with the protein-coding region of *RPSA*. No copy number variations were identified (see Methods). The overall proportion of ICA kindreds carrying protein-coding mutations of *RPSA* (12 of the 33 new kindreds) is consistent with both our previous report (8 of the 23 previous kindreds) and our model for the genetic architecture of *RPSA* mutations underlying ICA.

### The ICA cohort is enriched in *RPSA* protein-coding mutations

We then tested the hypothesis that these very rare nonsynonymous mutations of *RPSA* caused ICA, by comparing their frequency in the ICA cohort and the general population (15). The gnomAD and Bravo databases were considered the most suitable for such an *RPSA*-burden test as, together, they contain sequencing information for about 375,000 chromosomes, and the exome and genome sequences they contain provide good coverage of the protein-coding region of *RPSA*. We restricted our search to mutations of *RPSA* absent from the combined gnomAD and Bravo data, based on our previous estimates. Thus, for the calculation of *RPSA* burden, we considered only mutations that passed the quality filter and were present on no more than one chromosome in gnomAD and Bravo for our original comparison. We did not restrict our analysis to a specific population, as our cohort is diverse and the variants compared are extremely rare. We found that the newly recruited ICA kindreds, and the entire ICA cohort, were significantly enriched in very rare nonsynonymous variants, relative to the general population (*p*=2.44×10^−31^ and *p*=9.95×10^−51^ respectively) (**Table 1**). Restriction of the analysis to very rare missense variants and very rare in-frame indels revealed the newly recruited ICA kindreds, and the entire ICA cohort, to be significantly enriched in these variants (*p*=9.89×10^−21^ and *p*=3.44×10^−35^, respectively) (**Table 1**). Finally, we compared the nonsense, frameshift, and essential splicing mutations (in coding exons) in the ICA cohort with those present in the public databases, regardless of frequency. On the 375,000 chromosomes of gnomAD and Bravo, only two essential splice variants and three rare potentially damaging frameshift or nonsense variants (p.W176*, p.E235Vfs*60 and p.Q283*) were identified in the canonical ENST00000301821 transcript of *RPSA* (burden test *p*=2.85×10^−19^) (**Table 1**, **Table S2**, **Table S3**, **Figure S4**). The end of the RPSA sequence is less strongly conserved than the rest, so the frameshift of the last 60 amino acids, and the p.Q283* mutation removing the last 12 amino acids may have little effect. Overall, these results confirm our original observation that very rare non-synonymous mutations of *RPSA* cause ICA in about a third of the ICA kindreds.

**Table 1.** Statistical analysis of non-synonymous variants from ICA cohort versus gnomAD and Bravo database

|  | gnomAD + BRAVO total (WES + WGS) | Probands in original ICA kindreds 2013 | Test* | Probands in new ICA kindreds 2017 | Test** | Probands in entire ICA cohort | Test* |
| --- | --- | --- | --- | --- | --- | --- | --- |
| Total # of chromosomes in group | 375,000 | 46 |  | 66 |  | 112 |  |
| # of chromosomes with a very rare non-synonymous variant<sup>&</sup> | **86** | 8 | $2.97 \times 10^{-21}$ | **12** | **$2.44 \times 10^{-31}$** | **20** | **$9.95 \times 10^{-51}$** |
| # of chromosomes with a very rare missense variant, or inframe indel | 67 | 6 | $4.10 \times 10^{-16}$ | 8 | $9.89 \times 10^{-21}$ | 14 | $3.44 \times 10^{-35}$ |
| # of chromosomes with a nonsense, frameshift or essential splice variant<sup>#</sup> | 5 | 2 | $3.09 \times 10^{-07}$ | 4 | $1.10 \times 10^{-13}$ | 6 | $2.85 \times 10^{-19}$ |

\*: statistical significance threshold after correction with the stringent Bonferroni correction: $2.5 \times 10^{-6}$ (in our 2013 study, we looked at 4,222 genes and therefore we originally used a p-value cutoff of $1.2 \times 10^{-05}$ )
\*\*: statistical significance threshold: 0.05.
<sup>&</sup>: Non-synonymous variants were defined as nonsense, stop-loss, frameshift indels, inframe indels, missense, essential splice and splice variants affecting exons 2 to 7. Splice variants affecting only exon 1 were not taken into account for the calculation concerning non-synonymous variants, because exon 1 is non-coding, but they were analyzed in the UTR burden test (**Table 2**). All of the variants used in the gene-burden tests are presented in **Tables S2 to S5**.

### Incomplete penetrance of some novel protein-coding RPSA mutations

Unlike that of the original *RPSA* protein-coding mutations reported in 8 ICA families (5), the penetrance for ICA of the mutations in 6 of the 12 new kindreds (50%) with new protein-coding mutations of *RPSA* was incomplete (**Figure 1C**). Fifteen individuals carrying an ICA-causing mutation were found to have structurally and functionally normal spleens based on abdominal US or CT scans and/or the absence of Howell-Jolly bodies on blood smears. The distribution of these 15 asymptomatic carriers of RPSA mutation was as follows: p.A21P (1 of 3 carriers), p.G26S (1 of 3), p.M34V (1 of 2), p.[I65I;V66F] (9 of 11), p.Q84R (1 of 2), and p.V197Sfs*26 (2 of 3). We hypothesized that the nature of the *RPSA* mutation (e.g. its biochemical impact) contributes to the penetrance of ICA. We compared the Combined Annotation Dependent Depletion (CADD) scores of the mutations with their penetrance for ICA. CADD scores take into account information about evolutionary conservation, gene regulation, and transcription, to calculate the extent to which a nucleotide change is likely to be deleterious (see Methods) (16). There was no significant difference between the mean CADD of incompletely penetrant compared with fully penetrant *RPSA* mutations (mean CADD 25.8 vs. 29.4, p-value=0.58 (**Figure 2A**, **Figure S4**). Similarly, other deleteriousness predictors did not show significant differences between the two sets of variants (**Figure S5**). We then mapped positions of the human mutations onto the human crystal structure of the small ribosomal subunit (**Figure 2B**) (17). All five missense mutations with incomplete penetrance were found to be located outside of the hydrophobic core of RPSA lined with two α-helices (**Figure 2C, 2E**). Amino acid 180, a mutational hotspot in ICA modified in 5 of 14 families with completely penetrant *RPSA* mutations, contacts the nearby α-helix and seems to stabilize the core (**Figure 2C, 2D**). Moreover, all seven missense mutations with complete penetrance caused a change in the polarity or charge of the amino-acid side-chain group, whereas p.M34V and p.V66F did not. These findings suggest that the incomplete penetrance of nonsynonymous mutations of *RPSA* may be due to the residual function of the protein, but further studies are required to understand the difference in penetrance of different *RPSA* mutations.

**Figure 2:**
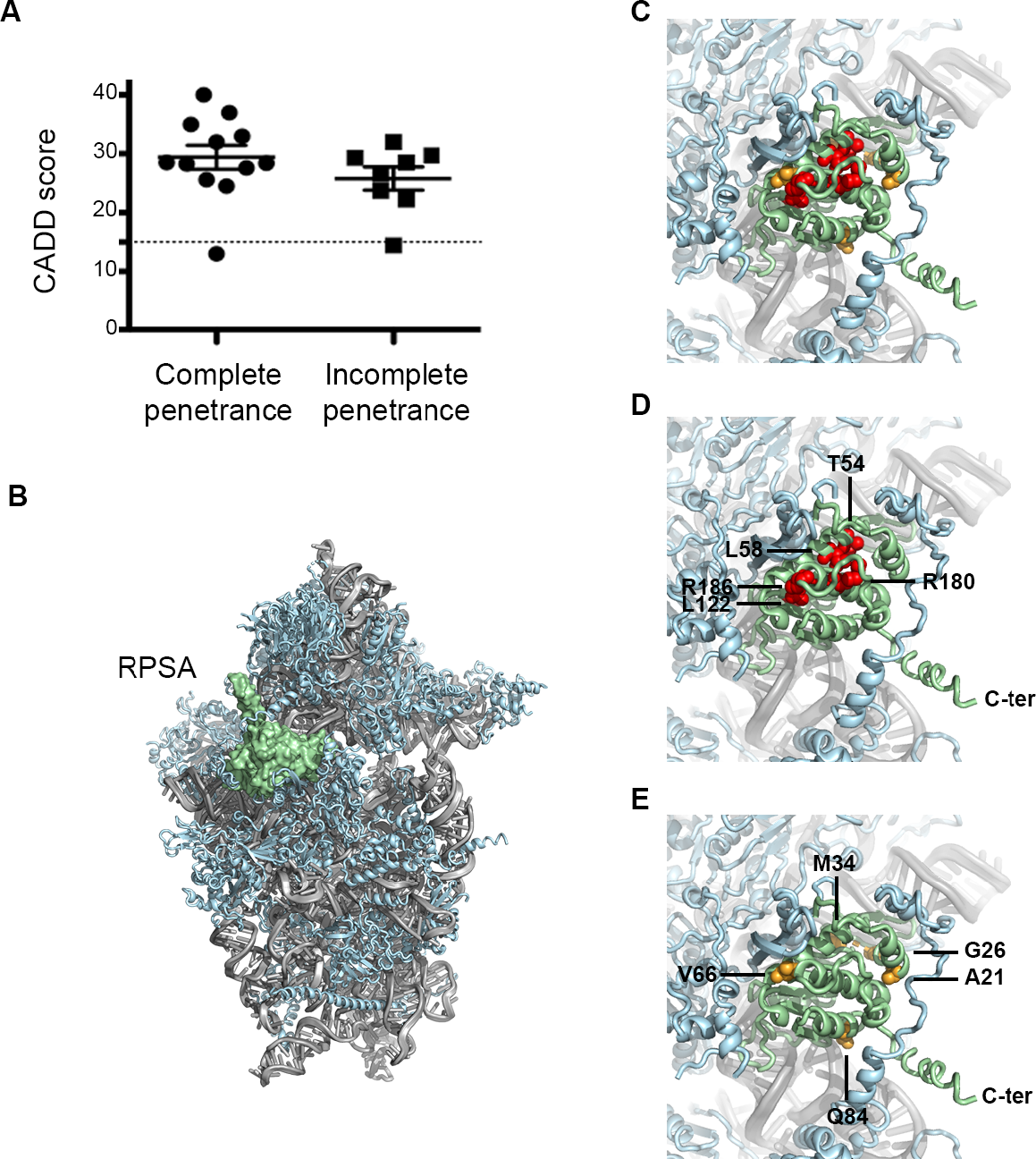
Several mutations in *RPSA* display incomplete penetrance for ICA. **A.**Comparative CADD score between RPSA coding-mutations leading to complete or incomplete penetrance. **B.**Structure of the human small ribosomal subunit (PDB 6EK0). 18S ribosomal RNA is colored in grey, ribosomal proteins are colored in light blue and RPSA is highlighted in green **C, D, E.**Detailed view of RPSA with colored spheres highlighting RPSA missense mutations leading to complete (red spheres) or incomplete (orange spheres) penetrance. Each amino acid found mutated in the ICA-cohort is indicated based on human RPSA.

### Identification of mutations in the non-protein-coding region of *RPSA*

We hypothesized that some cases of ICA may result from mutations in the non-coding region of *RPSA* exons. We searched for mutations in the 5’-UTR and 3’-UTR of *RPSA*, which are poorly covered by most WES kits. The 5’-and 3’-UTRs were subjected to Sanger sequencing in all 36 ICA kindreds of our cohort without very rare nonsynonymous mutations of *RPSA*. As for the candidate ICA-causing protein-coding mutations, we considered only mutations not present in gnomAD and Bravo. We identified two mutations predicted to affect the splicing of the 5’-UTR of *RPSA* that were private to our ICA cohort in three kindreds (**Figure 1D**). There were no very rare or private 3’-UTR mutations. One mutation, c.-34+5G>C (canonical ENST00000301821 transcript), was recurrent. It was identified in both a multiplex kindred (ICA-G) and a patient with sporadic disease (ICA-BD). Another mutation, c.-39_-34+3del (ENST00000301821), was identified in a patient with sporadic disease (ICA-AO). These two mutations were absent from the 150,000 chromosomes sequenced by WGS in gnomAD and Bravo corresponding to the subset of samples with good coverage of exon 1 of *RPSA*. Both 5’-UTR splice mutations were heterozygous. As for some of the protein-coding mutations, penetrance was incomplete in ICA kindreds carrying a mutation in the 5’-UTR of *RPSA*: c.-34+5G>C (1 in 4 of the individuals in ICA-G, and 2 of 3 in ICA-BD had a normal spleen) and c.-39_-34+3del (2 of 3). Overall, 3 of 36 ICA kindreds carried rare noncoding mutations in *RPSA* exons, whereas the overall proportion of ICA kindreds carrying a rare coding or noncoding mutation in *RPSA* exons was 23 of 56.

### Enrichment of the ICA cohort in *RPSA* non-coding mutations

We tested the hypothesis that these very rare non-coding mutations of *RPSA* cause ICA, by comparing their frequency in the ICA cohort and the general population. In total, 26 chromosomes carried a very rare *RPSA* 5’-UTR variant (1 chromosome in gnomAD WGS and Bravo, **Table S4**, **Table S5**). Fisher’s exact tests indicated that very rare mutations in the 5’-UTR or 3’-UTR of *RPSA* were more frequent in the ICA cohort than in the general population (*p*=9.75×10^−06^; **Table 2**). These results suggest that the very rare noncoding mutations identified in ICA patients may cause ICA. We also identified a 5’-UTR variant (c.-47C>T) in three unrelated Western European families that was absent from the gnomAD database but present in Bravo, with an allele frequency of 3.9×10^−05^. This variant was not retained as potentially disease-causing in our study, due to its allele frequency, but it might have a modifying effect (see discussion). The two kindreds carrying the c.-34+5G>C mutation were both from the United Kingdom. This mutation is not present in the UK10K dataset (18) corresponding to high-quality WGS data for 3,781 samples. We investigated whether this very rare variant was recurrent due to a founder effect or a mutational hotspot. We defined the haplotypes encompassing this mutation in both kindreds. We identified an 850 kb region with a haplotype of 250 SNPs common to the two kindreds. The length of this common haplotype strongly suggests that the mutation was inherited through a founder effect in both kindreds. Using ESTIAGE (19), we estimated the most recent common ancestor (MRCA) to have occurred 163 generations ago (CI_95%_= 52-743). Assuming a generation time of 25 years, the MRCA was estimated to have lived 4,075 years ago (CI_95%_= 1300-18575). The ICA cohort is, thus, enriched in private 5’-UTR mutations, one of which is present in two distantly related families, with incomplete penetrance.

**Table 2.** Statistical analysis of UTR variants from ICA cohort versus gnomAD and Bravo database

|  | Bravo + gnomAD (WGS) | ICA Probands without non-synonymous RPSA mutation | Test* | Probands in entire ICA cohort | Test* |
| --- | --- | --- | --- | --- | --- |
| Total # of chromosomes in group | 150,000 | 72 |  | 112 |  |
| All very rare 5'-UTR mutations | 26 | 3 | $3.83 \times 10^{-7}$ | 3 | $1.45 \times 10^{-6}$ |
| All very rare UTR mutations | **51** | **3** | **$2.58 \times 10^{-6}$** | 3 | $9.75 \times 10^{-6}$ |

\*: In this analysis, the 5'-UTR and 3'-UTR of RPSA are studied because RPSA is a candidate gene based on the protein-coding mutations in the same gene. This type of mutations was not studied in our initial cohort in 2013. Therefore, statistical significance threshold is set at 0.05. Statistical significance threshold after correction with the stringent Bonferroni correction is: $2.5 \times 10^{-6}$ .

### Functional impact of the *RPSA* 5’-UTR mutations

We investigated whether these 5’-UTR mutations were responsible for ICA, by studying their functional impact. The 5’-UTR of *RPSA* is not generally well conserved (**Figure S5**). However, the two mutations found in our ICA cohort are located at positions chr3:39,406,769 (hg38) and chr3:39,406,759-39,406,767, which are highly conserved between 100 vertebrate species (**Figure S6**) (18). This suggests that the 5’-UTR mutations identified in ICA patients may have a functional impact (**Figure S4**). We then used computational tools to construct hypotheses concerning the potential functional impact of these 5’-UTR mutations. The c.-34+5G>C mutation was predicted to impair splicing at the exon1/intron1 junction by disrupting the acceptor site (method: NNsplice9.0 (20)). The c.-39_-34+3del mutation eliminates the exon1/intron1 splicing junction and would, therefore, also be predicted to impair the splicing of intron 1. NNsplice9.0 predicted that the next splicing site would be 70 base pairs further downstream. We also estimated the potential impact on splicing of the other 5’-UTR variants present in gnomAD or Bravo. Only two of these 52 variants were predicted to impair splicing, both disrupting the acceptor splicing site for the intron1/exon2 junction (**Table S6**). We then experimentally tested the hypothesis that these ICA-associated 5’-UTR mutations impair splicing. RNAseq analysis confirmed the predictions made: both the c.-34+5G>C and c.-39_-34+3del mutations resulted in the retention of the same 70 base pairs (67 bp in the case of c.-39_-34+3del given that 3 nucleotides are deleted) (**Figure 3A**). The two 5’-UTR mutations lead to a 50% wild-type:50% mutant allelic ratio (**Figure 3B**). These 70 or 67 extra nucleotides contain two potential ATG codons (**Figure S7**). We also performed TOPO-TA cloning on cDNAs generated from peripheral blood mononuclear cells (PBMCs) from kindred ICA-G (c.-34+5G>C) (**Figure S8**). Again, we found that the c.-34+5G>C mutation prevented correct splicing at the exon 1/intron 1 junction, creating an insertion of 70 bp at the end of exon 1. Both splice mutations of the 5’-UTR thus impair the structure but not the stability of the mRNA, contrasting with the completely penetrant p.D73Vfs*16 mutation, for which no *RPSA* transcript carrying the mutation (0/67) was identified by TOPO-TA cloning, likely due to nonsense-mediated decay. Overall, these results suggest that rare mutations with a significant impact on the 5’-UTR of *RPSA* probably cause ICA, with incomplete penetrance.

**Figure 3:**
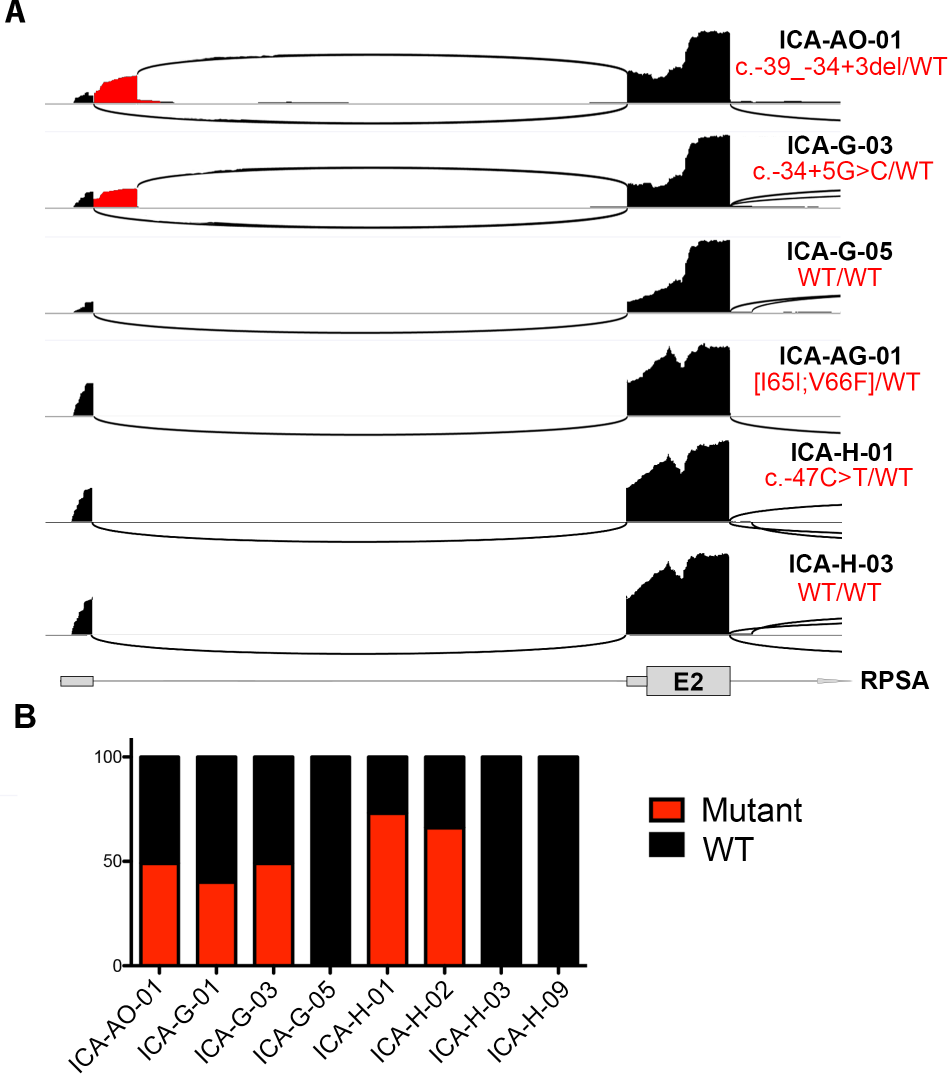
Impact of the two 5’-UTR mutations on *RPSA* mRNA structure. The two 5’-UTR mutations identified in ICA patients impair the splicing of intron 1 and lead to an intron retention of 70 or 67 bp at the end exon 1. **A.** Sashimi plots of RNAseq results for ICA-AO (c.-39_-34+3del); ICA-G (c.-34+5G>C) and ICA-H (variant c.-47 C>T). In ICA-AO and ICA-G the mutations lead to a 67 or 70 bp intronic retention respectively (red) whereas variant c.-47 C>T does not impact mRNA structure (ICA-H). **B.** Quantification of the allelic ratio. About half of the transcripts show the splicing defect in ICA-AO and ICA-G patients.

## Discussion

We identified eleven new protein-coding mutations and the first two non-coding mutations of *RPSA* responsible for ICA. Mutations in *RPSA* exons underlie ICA in 41% of index cases and kindreds (23 of 56 kindreds). This proportion differs significantly between multiplex kindreds (such mutations were found in 12 of 13 multiplex kindreds; 92%) and kindreds affected by sporadic ICA (11 of 43 kindreds; 24%). Mutations in *RPSA* exons were found to cause ICA in about half the patients (40 of 73) from the 56 kindreds included in our international ICA cohort. All of the mutations identified were not present in any variant database we looked at, which included more than 375,000 chromosomes. The burden of *de novo RPSA* mutations was high, with at least 7 of the 23 probands in kindreds with *RPSA* mutations carrying a *de novo RPSA* mutation. These results build on our previous report (5), as we have now identified mutations affecting both protein-coding and non-coding regions of the exons of *RPSA*. The genetic etiology of the ICA in kindreds without disease-causing mutations in the translated or untranslated regions of *RPSA* exons remains to be discovered. The relatively high frequency of *RPSA* mutations in this cohort suggests that other mutations may be present in regulatory regions outside of the exons of *RPSA* in some of these patients. A similar situation has been reported for blepharophimosis-ptosis-epicanthus inversus syndrome, for which causal mutations have been identified as much as 300 kb upstream from *FOXL2* (21).

An analysis of the relatives of ICA patients with *RPSA* mutations led to the identification of the first eight *RPSA* mutations with incomplete penetrance. This raises the question of the underlying cause for this inter-individual variability in phenotype. Mutations with incomplete penetrance are not predicted to have a lower functional impact, when compared with fully penetrant mutations, based on CADD or eleven other scores. As *RPSA* mutations cause ICA by haploinsufficiency, incomplete penetrance may be the result of hypomorphic mutations causing a smaller decrease of functional RPSA levels in heterozygotes than observed with loss-of-function mutations. The presence or absence of a modifier variant in a gene other than *RPSA*, or of an additional variant of *RPSA* (in *cis* or *trans*), rare or common, may also modulate the impact of the mutation (22). For example, a *RPSA* variant in *trans* to the mutation, increasing the levels of WT RPSA, may protect an individual heterozygous for a null or hypomorphic *RPSA* mutation. The analysis of the phenotypic variability between individuals carrying mutations in *RPSA* will either require a comprehensive analysis of very large kindreds, such as the ICA-AG kindred, or a sufficient level of genetic and perhaps allelic homogeneity across kindreds. An example of a common variant explaining the phenotypic variation in multiple individuals carrying a rare pathogenic variant is the digenic inheritance of non-syndromic midline craniosynostosis caused by rare *SMAD6* mutations and common *BMP2* variants (23).

The *RPSA* gene encodes ribosomal protein SA, a core component of the small subunit of the ribosome (24). There are 80 known human ribosomal proteins (25). AD ICA most likely results from haploinsufficiency, rather than negative dominance, because of the high proportion of truncating mutations and the strong purifying selection operating on the *RPSA* locus (5). Non-coding 5’-UTR ICA-causing mutations of *RPSA* provide further support for a mode of action based on haploinsufficiency. The molecular and cellular mechanisms by which RPSA haploinsufficiency underlies ICA have remained elusive. Even more intriguingly, mutations of the genes encoding 15 other human ribosomal proteins cause Diamond-Blackfan anemia (DBA), including mutations of *RPS19*, the most common genetic etiology of DBA (26). DBA (OMIM #105650) is a ribosomopathy underlying a syndromic form of aplastic anemia. Patients typically present with normochromic macrocytic anemia together with growth retardation, and about 30 to 50% have congenital craniofacial, upper limb, heart and/or urinary system malformations (27). None of the DBA patients reported in previous studies have asplenia, despite their wide range of developmental abnormalities. Conversely, none of the ICA patients apparently present any of the many hematological and developmental features of patients with DBA. The distinctive features of developmental tissues observed in the ribosomopathies underlying ICA and DBA, and emerging studies in other organisms suggest that the ribosome may influence the temporal and spatial control of gene expression during development. In an elegant study, lower amounts of ribosomes were shown to preferentially alter the translation of some transcripts, such as *GATA1*, thereby impairing erythroid lineage commitment and underlying DBA (28, 29).

Variable clinical expression, between patients and kindreds, of red-cell aplasia and congenital malformations has been widely reported for DBA (30, 31). The DBA-causing mutations of *RPS19* resulting in a severe clinical phenotype typically occur at a ‘mutation hot spot’ in the core of the protein, which has a structure consisting of five α-helix bundles organized around a central amphipathic α-helix (32). In RPSA, arginine in position 180 is a hotspot for ICA-causing mutations, and is also part of the core of the RPSA protein. We identified five completely penetrant mutations at the codon coding for R180. Incompletely penetrant mutations for both conditions map outside the core of the protein (**Figure 2C-E**). Moreover, the first exon of *RPS19*, like that of *RPSA*, is untranslated (5’-UTR) and the start codon is located at the start of exon 2. In addition to essential splice site mutations, one or two nucleotides downstream from the start codon of *RPS19* (33), rare 5’-UTR DBA-causing variants of *RPS19*, resulting in a longer 5’-UTR, have been shown to reduce RPS19 protein levels by 20%, modifying translational efficiency in a tissue-specific manner (34). The parallel between the *RPSA* mutations underlying ICA and the *RPS19* mutations underlying DBA is striking. In both cases, variants of both translated and untranslated regions of the exons can be disease-causing, and both types of mutations can show incomplete penetrance.

## Methods

### Patients

The publication of our paper identifying RPSA variants as a genetic etiology of ICA led to additional referrals. Nine families, ICA-AG, ICA-AL, ICA-AM, ICA-AO, ICA-AQ, ICA-AR, ICA-BH, ICA-BP and ICA-BO, approached us directly. Patients were recruited on the basis of three diagnostic criteria for ICA: (i) absence of a spleen on ultrasound or computed tomography, or absence/presence of a splenic remnant on autopsy, (ii) presence of Howell-Jolly bodies on blood smear, and (iii) absence of heart abnormalities. After weighing up the pros and cons, we decided not to consider the members of ICA families with an accessory spleen as ICA patients. On the one hand, it seems plausible that a mutation leading to ICA may affect normal spleen development and results in the development of an accessory spleen. This would provide an argument for both individuals with no spleen and those with an accessory spleen being included in the ‘case’ group for familial segregation analyses. On the other hand, accessory spleens are present in about 10% of the population, a frequency several orders of magnitude greater than that of total asplenia (35). As we have more than 50 kindreds enrolled in our cohort, we would expect there to be at least a few kindreds including individuals with an accessory spleen. We therefore decided not to consider these individuals as cases during familial segregation studies, and the absence of an *RPSA* mutation in these individuals would therefore not rule out the involvement of the mutation in ICA in other family members. In addition to studying the newly recruited families, we also updated the information for families reported in 2013. First, a newborn in family A was diagnosed with ICA. This baby carries the same missense mutation, p.R180G, as his relatives with ICA. Second, one patient, ICA-M-01, wild-type for *RPSA* has recently been diagnosed with Fanconi anemia, and we have identified compound heterozygous mutations of *FANCA*. Third, a sporadic boy (ICA-AH) has been found to carry a mutation in *ATRX*, which has been associated with asplenia as part of a broader syndrome in the past. He also suffers from mental retardation. We have retained these two kindred in all of the calculations in this paper to ensure consistency with our first report. Finally, the reader may note several gaps in the ‘numbering’ of the kindreds. For example, there is no kindred ICA-BA or ICA-BB, but there is an ICA-BC kindred. These gaps correspond to families or patients that have contacted us but that were not included in this study, for diverse reasons such as a lack of availability of DNA samples or of a signed informed consent form.

### Sanger sequencing of the seven exons of *RPSA*

In addition to the primers used for sequencing exons 2-7, as described in our previous paper (5), we designed primers to sequence the 5’-UTR/exon 1 of *RPSA*. We also included an M13 sequence in all primers (exons 1-7), to decrease the sequencing time. Primer-BLAST (www.ncbi.nlm.nih.gov/tools/primer-blast) was used to check that the primers would not amplify another sequence from the human genome. Following the isolation of genomic DNA from EBV-B cells or PBMCs from patients and their relatives, each of the seven exons of *RPSA* was sequenced with these primers (listed below).

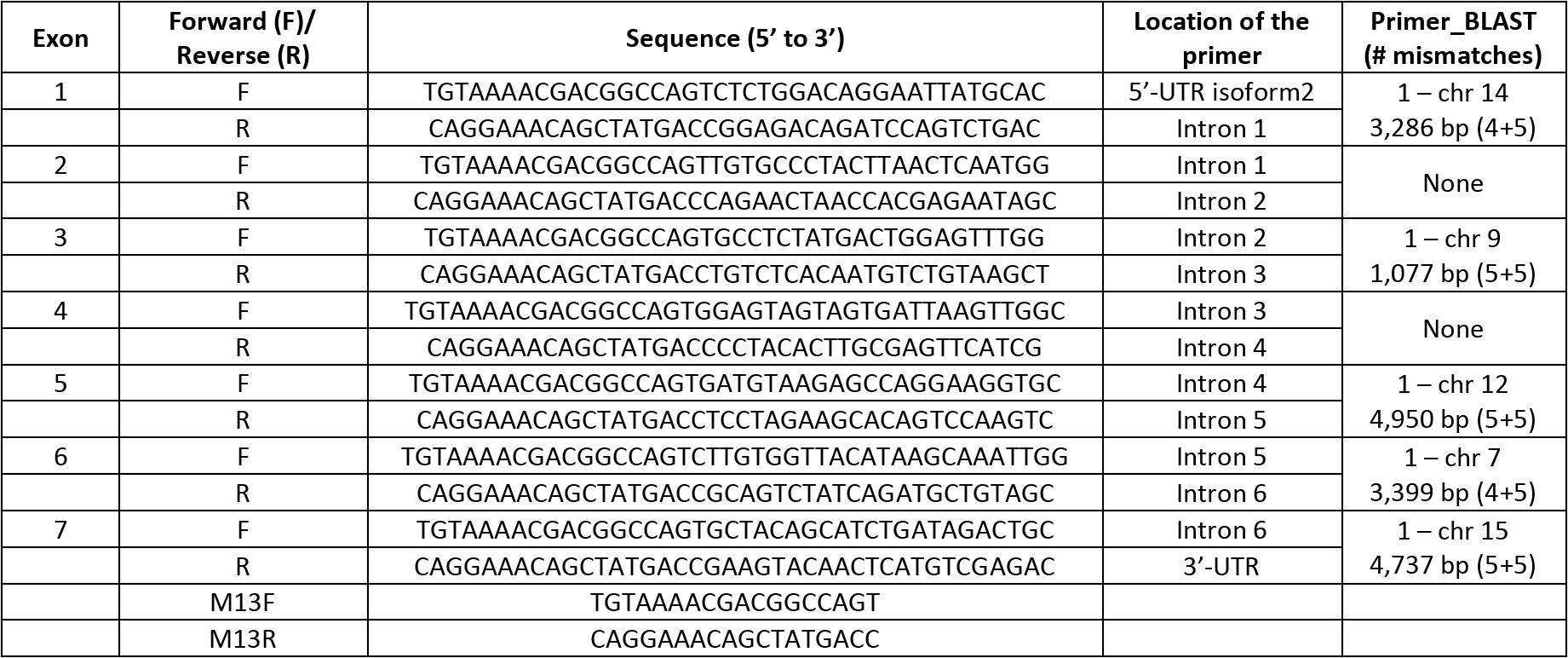

### Allele frequencies

The allele frequencies of *RPSA* variants present in the general population were retrieved from the exome and genome information in two public databases (last accessed February 2018). The gnomAD database contains 123,136 exome sequences and 15,496 whole-genome sequences from unrelated individuals sequenced as part of various disease-specific and population genetic studies. Information about *RPSA* can be obtained from the following URL: http://gnomad.broadinstitute.org/gene/ENSG00000168028 and the Bravo database lists genomic information for 62,784 possibly related individuals sequenced as part of NHLBI’s TOPMed program. Information for *RPSA* can be obtained from https://bravo.sph.umich.edu/freeze5/hg38/gene/ENSG00000168028. We also interrogated the UK10K dataset (https://www.uk10k.org/data.html) to determine whether the variant identified in two kindreds from the UK was present in the general population of the UK. The UK10K dataset contains 3,781 low-depth whole-genome sequences and aims to characterize genetic variation exhaustively, down to a minor allele frequency of 0.1% in the British population. We also sequenced *RPSA* in 29 Japanese controls from the *Centre de l’Etude de Polymorphisme Humain* (CEPH) panel because of the Japanese origin of one side of the ICA-AG family. None of the 29 Japanese controls carried the p.[I65I;V66F] mutation.

### Estimation of AF for ICA

Maximum tolerated reference allele counts were calculated with the Frequency Filter available from URL: https://cardiodb.org/allelefrequencyapp/. Based on the findings published in our first report, we predicted that ICA-RPSA would display monogenic inheritance, an incidence of about 1/600,000, an allelic heterozygosity of 0.25, a genetic heterozygosity of 0.3 and a penetrance of 0.75. Confidence was set at 0.95 and the reference population at 400,000 alleles.

### Gene burden tests

All the data from public databases was downloaded on 20 February 2018. All variants with an impact on the canonical transcript ENST00000301821 were analyzed. Bravo and gnomAD differ in their representation of ENST00000301821, particularly at the end of exon 3. We used the gnomAD representation, which is identical to the Ensembl description. We also used the Liftover tool (https://genome.ucsc.edu/cgi-bin/hgLiftOver) to facilitate comparisons between the two databases. We converted the GnomAD hg19 coordinates into hg38 coordinates, and performed manual checks to ensure that this process did not introduce any errors.

Based on our model, we focused principally on variants present on only one chromosome in the public databases. However, about 3,000 whole-genome sequences are present in both gnomAD and Bravo. We therefore decided to retain a variant that was present on one chromosome in Bravo and one chromosome in gnomAD. However, we counted it as occurring on only one chromosome in this case (1+1=1). Other variants present on two chromosomes in either database were systematically filtered out (2+0=0).

Nonsynonymous variants were defined as nonsense, stop-loss, frameshift, missense, essential splice and splice variants affecting exons 2 to 7. Splice variants affecting only exon 1 were not taken into account for the calculation concerning non-synonymous variants, because exon 1 is non-coding, but they were analyzed in the UTR burden test. Moreover, the whole-exome sequences of the gnomAD database do not cover this region. The c.*3_25delTGTTCTTGCATAGGCTCTTAAGC mutation was not taken into account for the loss-of-function variant burden test, as the deletion results in the same 295 amino acids and termination codon. All of the variants used in the gene-burden tests are presented in supplementary tables (**Tables S2 to S5**).

We restricted our RPSA-burden tests to variants that passed the quality filter in gnomAD and Bravo, as most of the low-quality variants are false positives (36). Fisher exact tests were performed with R software:

*input2.df = matrix(c(a,b,c,d), nrow = 2)*
*p_val = fisher.test(input2.df)$p.value*.

### Multiplex ligation-dependent probe amplification (MLPA) for CNV detection

We also searched for losses of one copy of *RPSA* as a cause of haploinsufficiency. We used the multiplex ligation-dependent probe amplification (MLPA) method, which has proved successful for the detection of CNVs in DBA (37). We designed probes mapping at least partly to the intronic region, because *RPSA* has several pseudogenes. As controls, we used probes binding to chromosomes 18, X and Y. We also designed a probe binding upstream from exon 1 to cover a region including a 21 bp deletion SNP (rs199844419) that would impair the binding of the probe, as a proxy for a control for the loss of one copy of *RPSA*. Three individuals in our cohort are heterozygous carriers of this SNP. No kindred presented a loss of one copy of one *RPSA* probe except for the probe upstream of exon 1 in the individuals carrying the 21bp deletion. We also searched the ExAC database (http://exac.broadinstitute.org/gene/ENSG00000168028) for the presence of CNVs in *RPSA*: none have been reported to date (last accessed January 2018). At the time of submission (March 2018), CNVs have not yet been included in the gnomAD or Bravo databases.

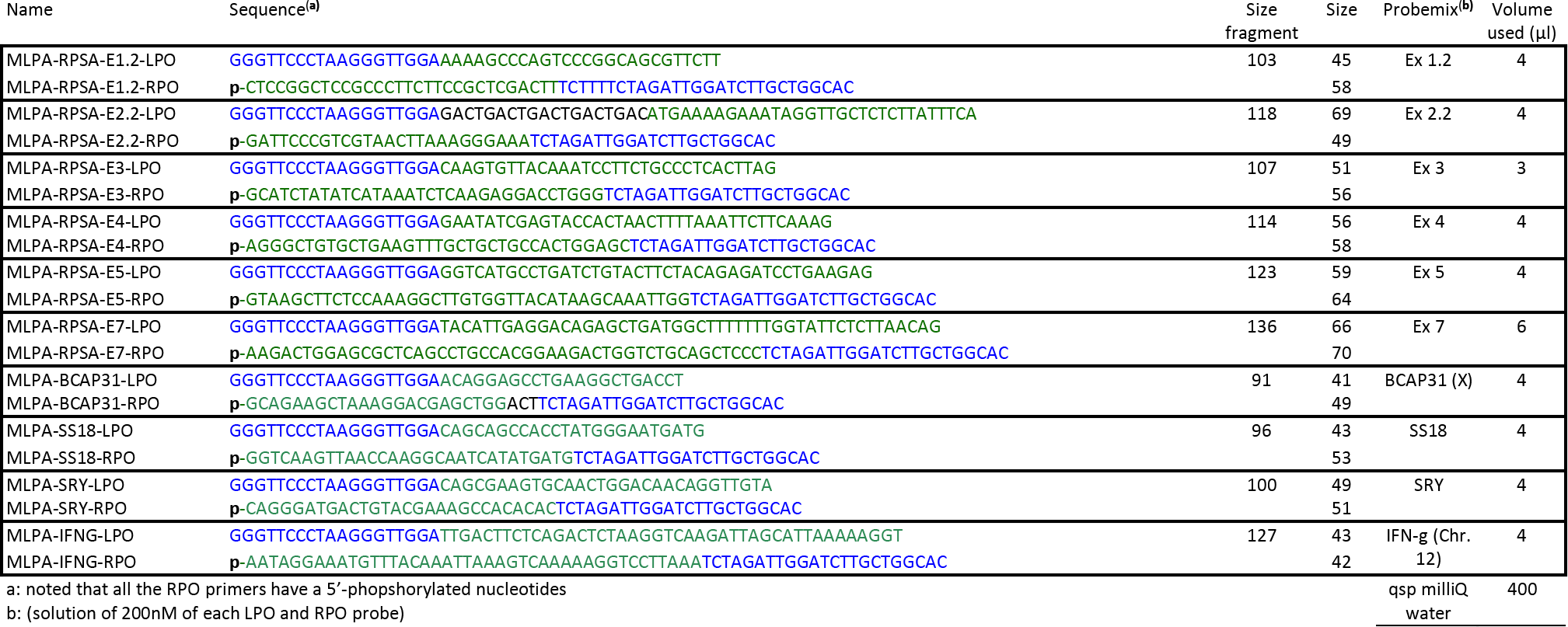

The MLPA protocol was performed using the reagent from MRC-Holland.

- The left and right probes (LPO and RPO) are mixed together to a final concentration of 200nM each in 1x Tris-EDTA (10mM Tris pH8.0, 1mM EDTA).
- All of the probes are then mixed and diluted using the volumes in the table below to prepare the Synthetic Basic Probemix.
- gDNA from individuals to be studied (100-150ng) was resuspended in a total of 5μl TE, and denatured (98°C, 5’; 25 °C, 5’).
- Synthetic Basic Probemix (1.5μl) and MLPA buffer (1.5μl) were added to the denatured gDNA and incubated at 95°C for 1’ followed by 60°C for 16-20 hours for hybridization. A 32μl of ligase mix buffer was then added to the mix for a 15’ incubation at 54°C, followed by an inactivation step (98°C, 5’). Amplification of the ligated products is performed by adding 10 μl of PCR mix (SALSA PCR) followed by PCR amplification for 35 cycles (95°C, 30”; (60°C, 30”, 72°C, 1’) x35; 72°C, 20’; 15°C).
- The separation of fragments by capillary electrophoresis were performed mixing 1μl of undiluted PCR product to 9.0μl formamide and 0.2μl LIZ standard, and then denatured at 85°C for 2’ and hold at 4°C for a minimum of 5’ before analyzing using a ABI-3730 DNA Analyzer.

### *RPSA* haplotypes investigated by microsatellite analysis (*de novo*)

We determined whether the samples sequenced in kindreds in which *RPSA* mutations appeared *de novo* were of the expected origin (i.e. the sample labeled as the father indeed corresponded to DNA from the father, and that labeled as the mother indeed corresponded to the DNA of the mother. In other words, we checked that the parent-child relationships between the subjects were as indicated by the family, and that there was no mixing up of the tubes at any point during the study). This was necessary to determine whether *RPSA* mutations truly occurred *de novo* in the ICA-AN, ICA-AT, ICA-AV, ICA-BH and ICA-BU families. We thus analyzed microsatellites throughout the genome for the samples sequenced (proband, father and mother). Microsatellite analysis was performed with the Identifiler Plus kit (Thermo Fisher Scientific^®^) according to the manufacturer’s protocol, optimized for use on an ABI Prism 3730 DNA Analyzer (Applied Biosystems^®^). PCR was first performed on a Veriti^®^ 96-well thermal cycler, with 10 μL AmpFlSTR Identifiler Plus Master Mix, 5 μL AmpFlSTR Identifiler Plus Primer Set and 10 μL of a 0.1 ng/μL solution of DNA from the subject to be tested. The samples were then prepared for electrophoresis, using 1 μL of the PCR products or an Allelic ladder and 9 μL of a 1:9 mixture of GeneScanTM 600 LIZ^®^ Size Standard and Hi-Di Formamide. Finally, data were analyzed with Gene mapper 4.0. The results are shown in **Figure S3**.

### Haplotype analysis for c.-34+5G>C

SNPs were genotyped with an Affymetrix 6.0 chip. After quality control (including a call rate of 100%), 909,622 SNPs were retained. Genotypes were phased with SHAPEIT v2 (38) software, with the 1000 Genomes samples as a reference panel. Familial information was used to improve the phasing and to identify the paternal haplotypes in which the mutation was located. We then determined the shared haplotype around the mutation and applied the likelihood-based ESTIAGE method (19) to estimate the age of the most recent common ancestor (MRCA) for the mutation. Recombination rates and haplotype frequencies were provided by the HapMap Project.

### Prediction of the impact of 5’UTR mutations on mRNA structure

We used the NNsplice 9.0 tool (http://www.fruitfly.org/cgi-bin/seq_tools/splice.pl) (20) to assess the potential impact on splicing of the rare variants at the exon1/intron1 splice junction identified in three kindreds with ICA. This tool has been shown to perform well for predicting the impact of variants leading to a splicing defect (https://github.com/counsylresearch/posters/raw/gh-pages/2017/ASHG/Q506C_Tran_Intronic_Variants_V1_Final_PRINT.pdf).

### RPSA structure and modeling of the mutations

The structural data were obtained by mapping the positions of the human mutations onto the human crystal structure of the small ribosomal subunit (Entry #6EK0) (17). All structural analyses and illustrations were made using PyMOL Molecular Graphics System, v.1.8 (Schrödinger).

### RNAseq

Total cellular RNA was isolated from PBMCs with the RNeasy Mini kit (QIAGEN^®^). Multiplexed RNA libraries were prepared with the Truseq RNA sample prep kit (Illumina^®^). In brief, poly(A)-containing mRNAs were captured with oligodT beads, fragmented, reverse-transcribed, and the cDNA was ligated to Illumina adapters containing indexing barcodes. Libraries were quantified with KAPA Library Quant kits (KAPA Biosystems), and run on a HiSeq 2000 Sequencing System (Illumina) to produce 150 bp single-end reads. Reads were then mapped with STAR v2.5.3a. Sashimi plots were generated with IGV (Broad Institute), using a minimum threshold of 20 reads for a splice event to be taken into account. The RNAseq dataset used will be available here: https://www.ncbi.nlm.nih.gov/bioproject/477701 on August 1, 2018 (biosamples: SAMN09476211, SAMN09476212, SAMN09476213, SAMN09476214, SAMN09476215, SAMN09476216)

### TOPO-TA cloning

RNA extracted from PBMCs was reverse-transcribed with the Superscript III First-Strand synthesis system for RT-PCR kit (18080-051), following the manufacturer’s protocol, with random hexamers. The TOPO TA cloning kit for sequencing (Invitrogen^®^#450030) was then used for subsequent cloning. The products were used to transduce Oneshot TOP 10 competent cells. Finally, we performed PCR on these colonies and sequenced them with the cRPSA-WT-I1-R /cRPSA-WT-R6 primers (exons 1 to 4).

## Acknowledgments

We thank the patients and all of their families. Many of the new families in this study contacted us directly and devoted considerable effort and time to their participation in this study. We also thank Yelena Nemirovskaya, Dominick Papandrea, David Hum, Cécile Patissier, Benedetta Bigio and Maya Chrabieh, who helped with all of the administrative questions and tasks required for this study. This work was supported in part by the March of Dimes (no. 1-FY12-440), St. Giles Foundation, National Center for Research Resources and the National Center for Advancing Sciences (NCATS) of the National Institutes of Health (NIH) (no. 8UL1 TR001866), the French National Agency for Research (ANR) under the “Investissement d’avenir” program (no. ANR-10-IAHU-01), and the Integrative Biology of Emerging Infectious Diseases Laboratory of Excellence (no. ANR-10-LABX-62-IBEID). A. Bol was funded by a fellowship from the Jane Coffin Childs Memorial Fund for Medical Research from July 2014 to November 2015. B. Bos. was supported by a fellowship from the Boehringer Ingelheim Fonds.

## Author contributions

A.Bol. conceived and designed the study. A. Bol., B. Boi., B. Bos., A.A., M.B., P.S., C-M.M. performed generated and analyzed the data. V.B., T.B., E.C., A.E., A.F., F.G., A.H., M.J., J.K., R.K., C-L.K., D.K., S.L., A.L-U., N.M., S.M., J.M.-G., M.N., M.O., M.P., C.P., A.J.P, C.R.G., C.T., H.V.-B., A.W., and I.M., recruited the patients, and provided clinical information. A. Bol., Boi., B. Bos., M.B., C-M.M., M.R., L.S., A.P., S.K., L.A., J.-L.C. interpreted the data. A. Bol., B. Boi., B. Bos., J.-L.C. wrote the manuscript. All authors reviewed the manuscript before submission.

## Abbreviations

AD: autosomal dominant
AF: allele frequency
CADD: Combined Annotation Dependent Depletion
CNV: copy number variation
DBA: Diamond-Blackfan anemia
ICA: isolated congenital asplenia
MLPA: multiplex ligation-dependent probe amplification
MRCA: most recent common ancestor
MSA: microsatellite analysis
OMIM: Online Mendelian Inheritance in Man
RPSA: ribosomal protein SA
WES: whole-exome sequencing
WGS: whole-genome sequencing
US: ultrasound
UTR: untranslated region

## Prediction software

CADD: Kircher M, et al. (2014) Nat Genet 46(3):310–315 http://cadd.gs.washington.edu/download

Condel: Gonzalez-Perez A & Lopez-Bigas N (2011) Am J Hum Genet 88(4):440–449. http://bbglab.irbbarcelona.org/fannsdb/query/condel

Envision: Gray VE, et al. (2018) Cell Systems 6(1):116–124.e113. https://envision.gs.washington.edu/

FATHMM: Shihab HA, et al. (2013) Human mutation 34(1):57–65 http://fathmm.biocompute.org.uk/fathmmMKL.htm

Grantham: Grantham R (1974) Science 185(4154):862.

Mutation Assessor: Reva B, et al. (2011) Nucleic Acids Res 39(17):e118–e118 http://mutationassessor.org/r3/

MCAP: Jagadeesh KA, et al. (2016) Nature Genetics 48:1581. http://bejerano.stanford.edu/mcap/

MPC: Samocha KE, et al. (2017) bioRxiv. ftp.broadinstitute.org/pub/ExAC_release/release1/

Polyphen2: Adzhubei IA, et al. (2010) Nat Methods 7(4):248–249 http://genetics.bwh.harvard.edu/pph2/

Provean: Choi Y, et al. (2012) Plos One 7(10):e46688. http://provean.jcvi.org/genome_submit_2.php?species=human

Revel: Ioannidis NM, et al. (2016) Am J Hum Genet 99(4):877–885.https://sites.google.com/site/revelgenomics/downloads

Sift: Kumar P, Henikoff S, & Ng PC (2009) Nature protocols 4(7):1073–1081. http://sift.jcvi.org/

## References

1. Myerson RM & Koelle WA (1956) Congenital absence of the spleen in an adult; report of a case associated with recurrent Waterhouse-Friderichsen syndrome. N Engl J Med 254(24):1131–1132.

2. Mahlaoui N, et al. (2011) Isolated congenital asplenia: a French nationwide retrospective survey of 20 cases. The Journal of pediatrics 158(1):142–148, 148 e141.

3. Maggadottir SM & Sullivan KE (2013) The diverse clinical features of chromosome 22q11.2 deletion syndrome (DiGeorge syndrome). The journal of allergy and clinical immunology. In practice 1(6):589–594.

4. Palamaro L, et al. (2014) FOXN1 in organ development and human diseases. International reviews of immunology 33(2):83–93.

5. Bolze A, et al. (2013) Ribosomal protein SA haploinsufficiency in humans with isolated congenital asplenia. Science 340(6135):976–978.

6. Thiruppathy K, Privitera A, Jain K, & Gupta S (2008) Congenital asplenia and group B streptococcus sepsis in the adult: case report and review of the literature. FEMS immunology and medical microbiology 53(3):437–439.

7. Gilbert B, et al. (2002) Familial isolated congenital asplenia: a rare, frequently hereditary dominant condition, often detected too late as a cause of overwhelming pneumococcal sepsis. Report of a new case and review of 31 others. Eur J Pediatr 161(7):368–372.

8. Takahashi F, et al. (2008) Isolated congenital spleen agenesis: A rare cause of chronic thromboembolic pulmonary hypertension in an adult. Respirology 13(6):913–915.

9. Rose C, et al. (1993) [Congenital asplenia, a differential diagnosis of essential thrombocythemia]. Presse medicale 22(34):1748.

10. Ferlicot S, Emile JF, Le Bris JL, Cheron G, & Brousse N (1997) [Congenital asplenia. A childhood immune deficit often detected too late]. Ann Pathol 17(1):44–46.

11. Lek M, et al. (2016) Analysis of protein-coding genetic variation in 60,706 humans. Nature 536(7616):285–291.

12. Rieux-Laucat F & Casanova JL (2014) Immunology. Autoimmunity by haploinsufficiency. Science 345(6204):1560–1561.

13. Boisson B, Quartier P, & Casanova JL (2015) Immunological loss-of-function due to genetic gain-of-function in humans: autosomal dominance of the third kind. Curr Opin Immunol 32:90–105.

14. Whiffin N, et al. (2017) Using high-resolution variant frequencies to empower clinical genome interpretation. Genet Med 19(10):1151–1158.

15. Lee S, Abecasis GR, Boehnke M, & Lin XH (2014) Rare-Variant Association Analysis: Study Designs and Statistical Tests. Am J Hum Genet 95(1):5–23.

16. Kircher M, et al. (2014) A general framework for estimating the relative pathogenicity of human genetic variants. Nat Genet 46(3):310–315.

17. Natchiar SK, Myasnikov AG, Kratzat H, Hazemann I, & Klaholz BP (2017) Visualization of chemical modifications in the human 80S ribosome structure. Nature 551(7681):472–477.

18. Pollard KS, Hubisz MJ, Rosenbloom KR, & Siepel A (2010) Detection of nonneutral substitution rates on mammalian phylogenies. Genome Res 20(1):110–121.

19. Genin E, Tullio-Pelet A, Begeot F, Lyonnet S, & Abel L (2004) Estimating the age of rare disease mutations: the example of Triple-A syndrome. J Med Genet 41(6):445–449.

20. Desmet FO, et al. (2009) Human Splicing Finder: an online bioinformatics tool to predict splicing signals. Nucleic Acids Res 37(9).

21. Crisponi L, et al. (2004) FOXL2 inactivation by a translocation 171 kb away: analysis of 500 kb of chromosome 3 for candidate long-range regulatory sequences. Genomics 83(5):757–764.

22. Castel SE, et al. (2017) Modified penetrance of coding variants by cis-regulatory variation shapes human traits. bioRxiv.

23. Timberlake AT, et al. (2016) Two locus inheritance of non-syndromic midline craniosynostosis via rare SMAD6 and common BMP2 alleles. eLife 5.

24. Malygin AA, Babaylova ES, Loktev VB, & Karpova GG (2011) A region in the C-terminal domain of ribosomal protein SA required for binding of SA to the human 40S ribosomal subunit. Biochimie 93(3):612–617.

25. Khatter H, Myasnikov AG, Natchiar SK, & Klaholz BP (2015) Structure of the human 80S ribosome. Nature 520(7549):640–645.

26. Mirabello L, et al. (2017) Novel and known ribosomal causes of Diamond-Blackfan anaemia identified through comprehensive genomic characterisation. J Med Genet 54(6):417–425.

27. Campagnoli MF, et al. (2008) RPS19 mutations in patients with Diamond-Blackfan anemia. Human mutation 29(7):911–920.

28. Ludwig LS, et al. (2014) Altered translation of GATA1 in Diamond-Blackfan anemia. Nat Med 20(7):748–753.

29. Khajuria RK, et al. (2018) Ribosome Levels Selectively Regulate Translation and Lineage Commitment in Human Hematopoiesis. Cell.

30. Landowski M, et al. (2013) Novel deletion of RPL15 identified by array-comparative genomic hybridization in Diamond-Blackfan anemia. Human genetics 132(11):1265–1274.

31. Narla A, et al. (2016) A novel pathogenic mutation in RPL11 identified in a patient diagnosed with diamond Blackfan anemia as a young adult. Blood cells, molecules & diseases 61:46–47.

32. Gregory LA, et al. (2007) Molecular basis of Diamond-Blackfan anemia: structure and function analysis of RPS19. Nucleic Acids Res 35(17):5913–5921.

33. Boria I, et al. (2010) The ribosomal basis of Diamond-Blackfan Anemia: mutation and database update. Human mutation 31(12):1269–1279.

34. Badhai J, Schuster J, Gidlof O, & Dahl N (2011) 5 ’ UTR Variants of Ribosomal Protein S19 Transcript Determine Translational Efficiency: Implications for Diamond-Blackfan Anemia and Tissue Variability. Plos One 6(3).

35. Vikse J, et al. (2017) The prevalence and morphometry of an accessory spleen: A meta-analysis and systematic review of 22,487 patients. Int J Surg 45:18–28.

36. Durtschi J, Margraf RL, Coonrod EM, Mallempati KC, & Voelkerding KV (2013) VarBin, a novel method for classifying true and false positive variants in NGS data. BMC Bioinformatics 14 Suppl 13:S2.

37. Quarello P, et al. (2008) Multiplex ligation-dependent probe amplification enhances molecular diagnosis of Diamond-Blackfan anemia due to RPS19 deficiency. Haematologica 93(11):1748–1750.

38. Delaneau O & Zagury JF (2012) Haplotype inference. Methods Mol Biol 888:177–196.

